# Hippocampal DNA methylation promotes memory persistence by facilitating systems consolidation and cortical engram stabilisation

**DOI:** 10.1101/2024.02.25.581942

**Authors:** Janina Kupke, Stefanos Loizou, C. Peter Bengtson, Carsten Sticht, Ana M.M. Oliveira

**Author notes:** Ana M.M. Oliveira. Department of Molecular and Cellular Cognition Research, Central Institute of Mental Health, Medical Faculty Mannheim, Heidelberg University, 68159 Mannheim, Germany.

## Abstract

The long-term stabilization of memory traces or engram involves the rapid formation of cortical engrams during encoding that mature functionally over time guided by the activity of the hippocampus. The molecular mechanisms that regulate this process remain largely unknown. Here, we found that hippocampal DNA methylation converts short-lasting into long-lasting memories by promoting systems consolidation and the stabilization of cortical engrams.

---

It is well accepted that consolidation of episodic memories occurs on different time scales after the learning phase. The initial steps of consolidation rely on synaptic-level changes in the hippocampus orchestrated by gene transcription and *de novo* protein synthesis. Once acquired, the storage of episodic memory is progressively more dependent on neocortical regions through a process termed systems consolidation^1–3^. Recent evidence shows that neocortical engrams, specifically in the medial prefrontal cortex (mPFC), are rapidly generated at the time of encoding but only become functionally mature as the time progresses^1,4,5^. The hippocampus plays a pivotal role in guiding the gradual maturation and stabilization of cortical engram cells^1,5,6^. However, molecular correlates of systems-level memory consolidation and engram stabilization are largely unknown. Epigenetic events are enduring regulatory mechanisms that have been implicated in memory formation^7^. DNA methyltransferases are required for synaptic plasticity^8,9^ and memory^10–15^ and DNA methylation changes take place during the formation of enduring forms of contextual fear memory, both in the hippocampus^16,17^ and cortex^16^. However, whether DNA methylation is causally linked to long-term storage of episodic memory and associated engram stability remains to be investigated. In this study, we evaluated the contribution of hippocampal DNA methylation to these processes.

To investigate whether hippocampal DNA methylation impacts memory persistence, we manipulated the levels of the DNA-methyltransferase 3a2 (Dnmt3a2) in the dorsal hippocampus (dHPC) of mice (Fig. 1a) and trained the mice in a contextual-fear-conditioning (CFC) paradigm. We used a training protocol that induces a fear memory that decays within 2 weeks^18^ (Fig. 1b). When comparing the group overexpressing Dnmt3a2 and the control, we observed no difference at recent memory recall (24h) (Fig. 1b). However, we found that overexpression of Dnmt3a2 converted the short-lasting into a persistent fear memory, as evidenced by high freezing rates 2 weeks after training (Fig. 1b). To determine whether this effect depends on the catalytic activity of Dnmt3a2, we tested the effect of overexpression of a catalytic inactive form of Dnmt3a2^19^ on remote memory recall (Extended Data Fig. 1a, b). In contrast to the overexpression of wild-type Dnmt3a2, the expression of the catalytic mutant had no effect on freezing behaviour during remote memory recall (Extended Data Fig. 1c), suggesting that the DNA methyltransferase activity is required for the conversion into long-lasting memory.

**Fig. 1.**
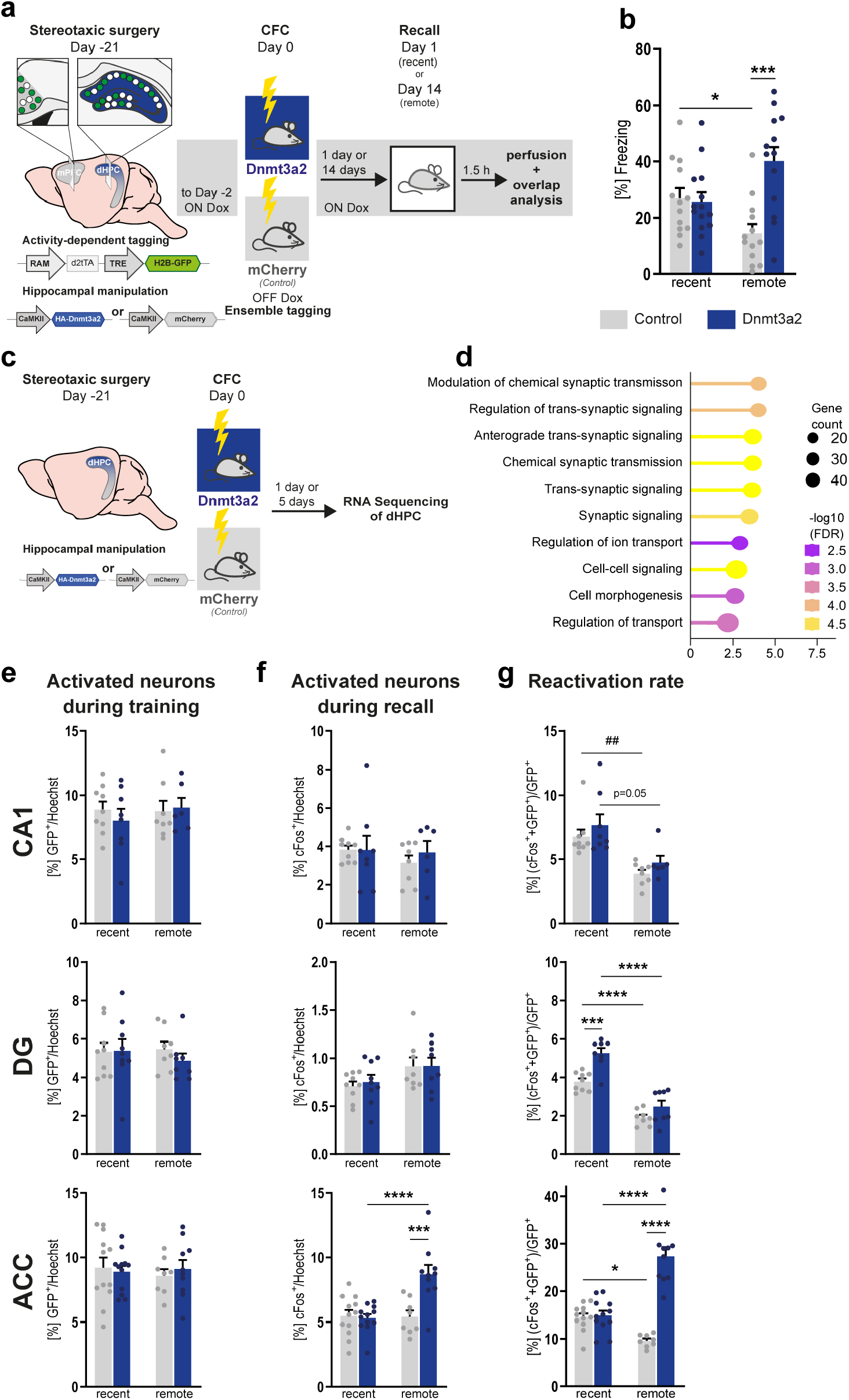
Hippocampal DNA methylation drives memory persistence and facilitates system consolidation. (a) Schematic representation of experimental design. Stereotaxic injection of rAAV to tag the CA1, DG and ACC memory engram and rAAV injection in dHPC to manipulate Dnmt3a2 levels. Three weeks after surgery, mice were trained in CFC (1x 0.2 mA shock) and memory was tested in a recall session 1day or 14 days after. 1.5h post recall, mice were sacrificed for overlap analysis. (b) Recent or remote long-term contextual fear memory of mice injected with rAAVs overexpressing mCherry (n=13 (recent); n=13 (remote)) or Dnmt3a2 (n=14 (recent); 13 (remote)). ***p≤0.001, *p<0.05 by one-way ANOVA test followed by Sidak’s multiple comparisons test. (c) Schematic representation of experimental design. Stereotaxic injection of rAAV in dHPC to manipulate Dnmt3a2 levels. Three weeks after surgery, mice were trained in CFC (1x 0.2 mA shock) and either 1 day or 5 days later, dHPC tissue was collected and RNA extracted for RNA Sequencing analysis. (d) GO-Term analysis of DEGs. Dot plot illustrates Top 10 GO term enrichment of biological processes. (e, f, g) Quantitative image analysis of CA1, DG and ACC region of (e) activated neurons during training assessed by GFP signal, (f) activated neurons during recall identified by endogenous cFos labelling, (g) reactivation rate (GFP^+^+cFos^+^ neurons in GFP^+^ population) (n=7-12). *p<0.05, ***p≤0.001, ****p≤0.0001, ns: not significant by one-way ANOVA test followed by Sidak’s multiple comparisons test. Error bars represents S.E.M.

To further address whether Dnmt3a2 overexpression impacts the hippocampal transcriptomic profile and infer possible functional consequences, we performed RNA-sequencing analysis of hippocampal tissue 1 or 5 days after CFC training (Fig. 1c). One day after CFC, 20 differentially expressed genes (DEGs) were found (Extended Data Fig. 1 d), which increased to 239 5 days after CFC (Extended Data Fig. 1 e) with an overlap of 13 genes (Extended Data Fig. 1 f). GO term analysis revealed a strong enrichment of functional terms related to synaptic regulation, e.g., “*Modulation of chemical synaptic transmission*”, “*Regulation of trans-synaptic signalling*”, and “*Synaptic signalling*” (Fig. 1d). We validated the RNA-sequencing results using independent biological samples by qRT-PCR and included home cage controls to see whether Dnmt3a2-overexpression causes changes in baseline conditions. We found several genes to be differentially expressed (Extended Data Fig. 1g), that were differentially methylated upon Dnmt3a2-overexpression in our previous study^15^, or *in vivo* after CFC^16,17^ (Extended Data Fig. 1h). Additionally, *Calb1* and *Fhad1* were differentially expressed upon Dnmt3a2-overexpression (Extended Data Fig. 1i) and increased in DG-engram neurons after CFC^20,21^. These findings indicate that the regulation of memory duration involves specific transcriptional changes in synaptic transmission-related genes that might be mediated by DNA methylation processes.

Next, we investigated whether the effect of Dnmt3a2 overexpression in memory duration impacts engram stability and maturation of supporting hippocampal and neocortical engram cells. In a first step, we established a tool that allows for the monitoring of engram dynamics in recent and remote memories. To this end, we virally tagged engram neurons in the dHPC and the anterior cingulate cortex (ACC) and trained mice in CFC paradigms that lead to either a strong, persistent fear memory or a short-lasting one (Extended Data Fig. 2a-c). The fraction of activated engram neurons during training (GFP^+^) was comparable across training protocols and time of testing, with exception of the dentate gyrus (DG) that showed a mild increase in strong versus weak protocols at remote time points. This may be attributed to variations in the tagging or viral injection (Extended Data Fig. 2d). These results suggest that the engram size does not depend on training strength, which is in line with previous studies ^22,23^. The analysis of neurons activated during memory recall (given by the percentage of cFos^+^ cells) revealed that in the dHPC, the proportion of activated neurons was similar during recent or remote recall, whereas in the ACC it increased during the retrieval of the strong long-lasting fear memory (Extended Data Fig. 2e). This finding is in line with previous studies and indicates an increased involvement of cortical regions during remote versus recent fear memory recall^3,24,25^. Next, we analysed the proportion of learning activated neurons that are reactivated during recall as a measure of engram reactivation (Extended Data Fig. 2f). At recent memory recall, the reactivation rate was similar across the different protocols and brain regions. At remote memory recall, engram reactivation decreased in all regions in mice trained with the short-lasting memory protocol (Extended Data Fig. 2f) in line with weak recollection of the fearful experience. In mice trained in the strong protocol, DG reactivation decreased, as expected due to the previously reported DG-independent role in remote memory recall^1,26^. In contrast, CA1 engram reactivation remained constant (Extended Data Fig. 2f) as shown by others^26,27^. As predicted, cortical engram reactivation increased over time (Extended Data Fig. 2f), in line with a functional engram maturation during systems consolidation^1,4,5^. After establishing the tool and confirming known engram dynamics during systems consolidation, we investigated the effect of hippocampal DNA methylation in this phenomenon. To this end, we combined Dnmt3a2-overexpression with engram tagging (Fig. 1a). Dnmt3a2-overexpression did not influence the size of the neuronal population activated by learning (GFP^+^) (Fig. 1e) nor did it cause changes in recall-activated neurons (Fos^+^) in DG or CA1 over time. However, overexpression of Dnmt3a2 in dHPC increased the number of recall-activated cells (Fos^+^) in ACC at remote memory recall (Fig. 1f). Engram reactivation decreased over time in all brain regions in the control group as well as in the dHPC in Dnmt3a2-overexpression conditions (Fig. 1g). Remarkably, in ACC, engram reactivation increased upon hippocampal Dnmt3a2-overexpression during remote memory recall (Fig. 1g). Taken together, these results show that hippocampal Dnmt3a2-overexpression converts a short-lasting memory into a persistent memory that shows characteristics typical of a naturally occurring remote fear memory (increased cortical engram reactivation and decreased hippocampal involvement). Thus, suggesting a role for hippocampal DNA methylation in cortical engram maturation and systems consolidation.

One characteristic of remote memory is that stored information loses contextual accuracy manifested by increased contextual generalisation compared to recent memory^28,29^. This has been attributed to the reduced involvement of the hippocampus during remote memory retrieval^30^. To further characterise the remote memory induced by Dnmt3a2 overexpression, we tested fear responses in an altered context at recent or remote times (Fig. 2a). We envisioned that if this manipulation facilitates the memory transfer from hippocampal towards cortical circuits, the resulting long-lasting memory will display reduced contextual specificity. At recent recall all mice showed low freezing behaviour in the altered context pointing towards contextual specificity (Fig. 2b). At remote recall, Dnmt3a2-overexpressing mice had higher freezing levels compared to recent time point (Fig. 2b). This demonstrates that the contextual information of the memory lost its specificity, which could indicate a cortical memory transfer. To further test whether the memory resides in cortical engram cells, we took advantage of the viral Targeted Recombination in Active Populations (TRAP) system^4^ that expresses the inhibitory Designer Receptors Exclusively Activated by Designer Drugs (DREADD) – hM4Di – only in cortical neurons activated during CFC training (Fig 2c). Mice received the viral TRAP in the ACC to express either hM4Di or mCherry (control) in engram cells and the Dnmt3a2-overexpression AAV in the dHPC (Fig. 2d). To test whether, in these conditions, cortical engram neurons are required for remote fear memory expression, the remote memory recall session was performed whilst inhibiting CFC-tagged hM4Di^+^ neurons (Fig. 2d). Inhibition of the ACC-engram by systemic administration of Clozapine-N-oxide (CNO) abolished remote fear memory retrieval (Fig. 2e), demonstrating that the memory trace resides in cortical circuits.

**Fig. 2.**
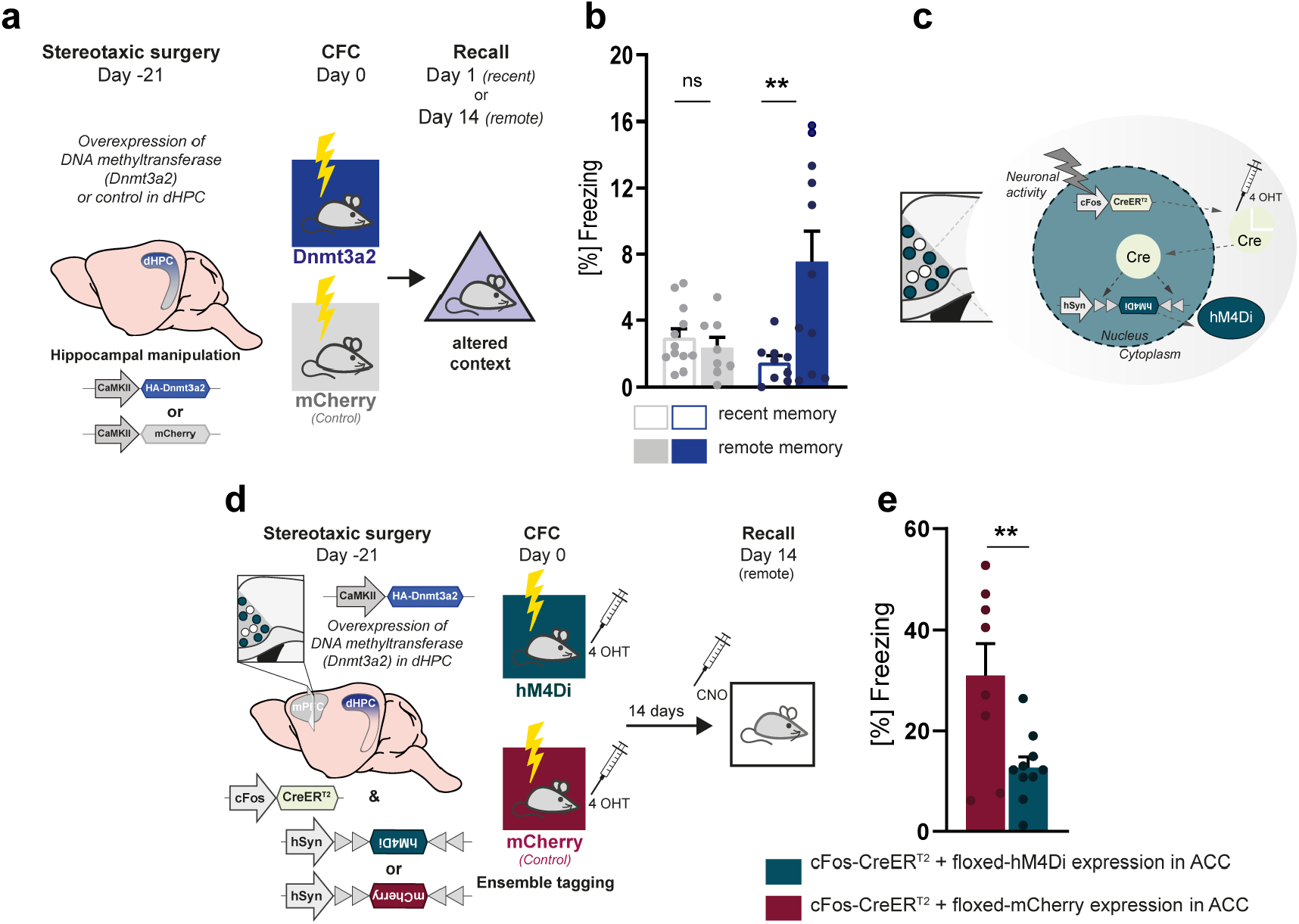
Hippocampal DNA methylation promotes the transfer of the memory trace to the cortex. (a) Schematic representation of experimental design. Stereotaxic injection of rAAV in dHPC to manipulate Dnmt3a2 levels. Three weeks after surgery, mice were trained in CFC and tested for their memory specificity in an altered context (either day 1 or 14). (b) Freezing behaviour in an altered context of mice trained in CFC and expressing mCherry (n=8-12) or Dnmt3a2 (n=9-11). **p≤0.01, ns: not significant by two-tailed, unpaired t-test. (c) Schematic representation of viral TRAP system. Neuronal activity triggers activation of the cFos promoter, which causes CreER^T2^ expression. By administering 4-hydroxytamoxifen (4-OHT) systemically, translocation of CreER^T2^ into the nucleus is allowed and irreversible recombination of the Cre-dependent vector is made possible. (d) Schematic representation of experimental design. Stereotaxic injection of rAAV to express Dnmt3a2 in dHPC and the viral TRAP in the ACC. Three weeks after surgery, mice were trained in CFC (1x 0.2 mA shock). To trap cortical engram neurons, 4-Hydroxytamoxifen (4 OHT) was injected *i*.*p*. 2h after CFC. 30 min prior memory recall session on day 14, mice were *i*.*p*. injected with Clozapine N-oxide (CNO) to inhibit the trapped cortical engram neurons and tested for remotememory. (e) Remote contextual fear memory of mice overexpressing Dnmt3a2 in dHPC and expressing viral trapped control construct (mCherry; n= 8) or the inhibitory DREADD (hM4Di; n=10) in ACC. **p≤0.01 by two-tailed, unpaired t-test. Error bars represents S.E.M.

We showed that the overexpression of Dnmt3a2 converts a short-lasting into a long-lasting fear memory that presents characteristics of a naturally occurring remote fear memory, supporting a function for hippocampal DNA methylation in remote memory formation and systems consolidation. To determine whether endogenous DNA methylation processes are crucial for memory persistence and systems consolidation, we assessed the effect of Dnmt3a2 depletion on memory duration using a CFC protocol that triggers remote memory and a validated shRNA construct^14^ (Fig. 3a, b). To exclude perturbation of memory encoding, we tested memory performance during recent memory retrieval (24h) and found the freezing rates not changed (Fig. 3c). Reduction of Dnmt3a2 led to remote memory impairments (Fig. 3d), implying that endogenously regulated DNA methylation processes are indeed crucial for memory duration. Next, we tested whether the remote memory impairment is associated with changes in cortical engram dynamics (Fig. 3e-g). While knockdown of hippocampal Dnmt3a2 did not affect the proportion of cells activated during training (Fig. 3e) or recall (Fig. 3f), it decreased cortical engram reactivation (Fig. 3g). Taken together, our loss-of-function experiments revealed that endogenous DNA methylation processes in the hippocampus are pivotal for memory strength and duration and regulation of cortical engram stabilisation.

**Fig. 3.**
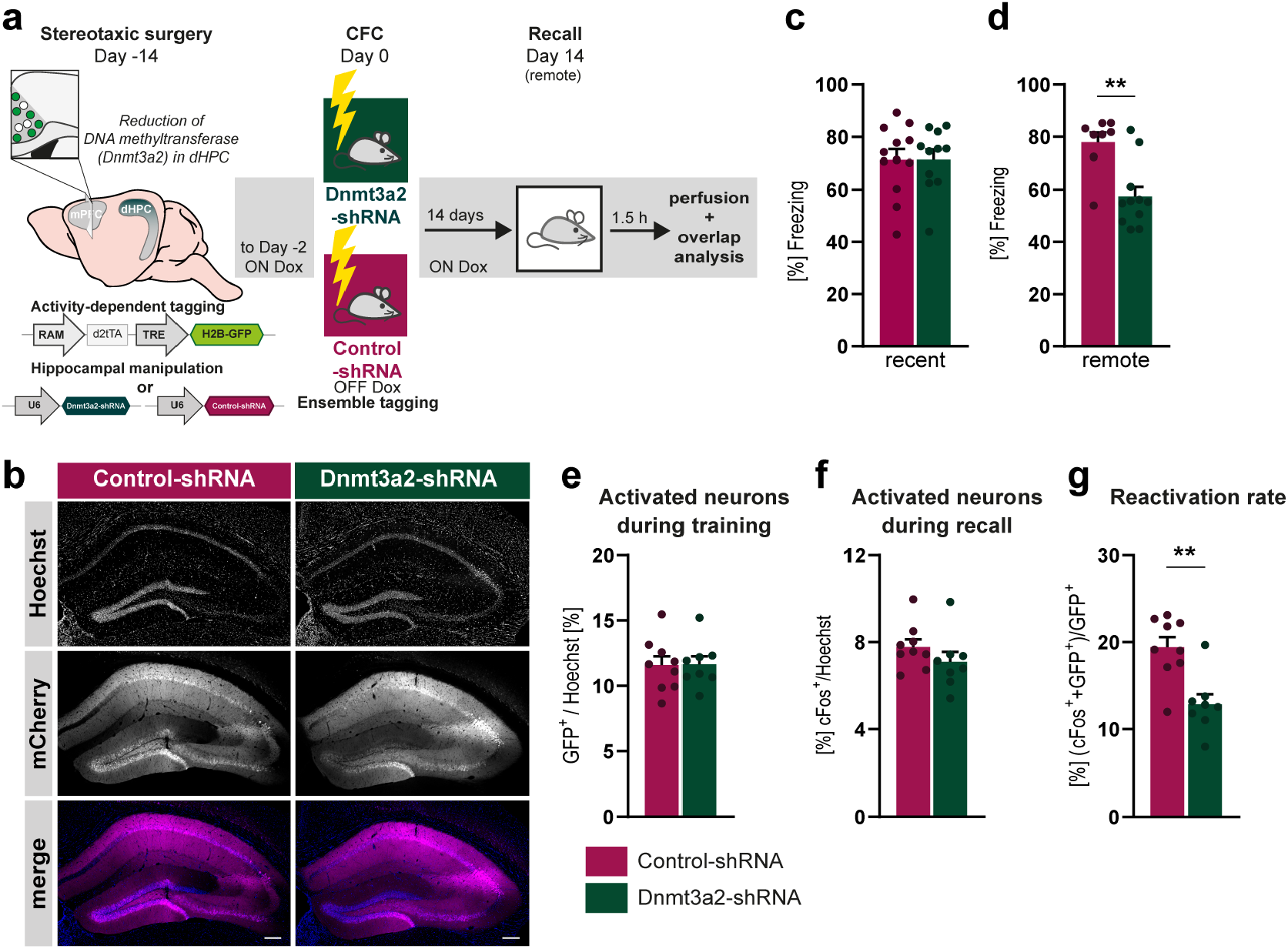
Endogenous DNA methylation processes regulate memory duration and cortical engram reactivation. (a) Schematic representation of experimental design. Stereotaxic injection of rAAV in the dHPC to knockdown Dnmt3a2 and into ACC to tag the cortical engram. Two weeks after surgery, mice were trained in CFC (1x 0.7 mA shock) and their memory was tested 14 days after. 1.5h post recall, mice were sacrificed for overlap analysis. (b) Representative images of dHPC of mice injected with rAAVs against Control-shRNA or Dnmt3a2-shRNA. Scale bar: 100μm. (c) Recent or (d) remote contextual fear memory of mice injected with rAAVs containing Control-shRNA (n=12(recent); n=8(remote)) or Dnmt3a2-shRNA (n=11 (recent); n=10(remote)). **p<0.01 by two-tailed, unpaired t-test. (e, f, g) Quantitative image analysis in ACC region of (e) activated neurons during training assessed by GFP signal, (f) activated neurons during recall identified by cFos labelling, (g) reactivation rate (GFP^+^+cFos^+^ neurons in GFP^+^ population) (n=8-9). **p≤0.01 by two-tailed, unpaired t-test. Error bars represents S.E.M.

In this study, we established a causal link between DNA methylation processes in the hippocampus and the long-term storage of memory. We showed that Dnmt3a2-overexpression in dHPC converted a short-lasting into a long-lasting memory while reducing Dnmt3a2 weakened a remote memory. Interestingly, increasing Dnmt3a2 did not impact hippocampal engram dynamics but increased cortical engram reactivation during remote memory recall, in line with a role in the maturation of cortical engram cells and systems consolidation^1,5^.

We found that Dnmt3a2 overexpression altered the expression of genes functionally associated with synaptic transmission and signalling. It is conceivable that DNA methylation-mediated gene expression changes modulate the communication between the hippocampus and neocortex that is required for systems consolidation^31,32^ and engram cell maturation^1,5^. Although the precise underlying mechanism remains to be defined, our study uncovered a previously unknown function for DNA methylation in fear memory engram maturation and stabilization.

## Methods

### Animals

3-month-old C57BL/6N male mice (Charles River, Sulzfeld, Germany) were used with *ad libitum* access to water and food, under a 12h light/dark cycle at 22 ± 1°C and 55 ± 10% relative humidity, and group-housed (2-3 mice per cage). Behavioural experiments were conducted during the light phase. Animals were randomly assigned to experimental groups and analysis was conducted blindly. All behavioural sessions were video recorded and manually scored to determine the freezing behaviour by an experimenter blind to the group identity. Procedures complied with German animal care guidelines and European Community Council Directive 86/609/EEC. After behaviour, histological analysis was performed to confirm correct targeting and tissue integrity. Mice that showed absence or missing viral expression were excluded.

### Recombinant Adeno Associated Virus (rAAV) production

Viral particles were produced and purified as described previously^18^. In brief, AAV-293 cells (Stratagene #240073, California, USA) were co-transfected with target AAV plasmid and helper plasmids (pFΔ6, pRV1, and pH21) using calcium phosphate precipitation. 60 hours after transfection, the HEK 293 cells were collected and lysed. Viral particles in media were concentrated by incubation with 40% Polyethylene glycol (PEG) solution (pH 7,4) for 3h at 4°C and then pelleted by centrifugation for 20 min. The pellet was added to lysed HEK cells prior to virus purification through heparin affinity columns (HiTrap Heparin HP, GE Healthcare, Uppsala, Sweden) and concentrated using Amicon Ultra-4 centrifugal filters (Millipore, Bedford, MA). The infection rate, toxicity, viral titer, and knockdown efficiency for each batch of generated viruses were evaluated. The final titer was around 1-2 ×10^12^ viral particles/ml. Expression of shRNAs was obtained by U6 promoter upstream of the shRNA sequence and a calcium-calmodulin kinase II promoter to drive mCherry expression. shRNA sequence was previously validated^33^. Sequences are as follows: Dnmt3a2-shRNA: ACGGGCAGCTATTTACAGAGC; Control-shRNA: ACTACCGTTGTTATAGGTG. For activity-dependent engram tagging, the H2BGFP sequence (H2B-GFP was a gift from Geoff Wahl (Addgene plasmid # 11680; http://n2t.net/addgene:11680; RRID: Addgene_11680) was subcloned into the pAAV-RAM-d2TTA::TRE-MCS-WPRE-pA construct (kindly deposited to Addgene by Dr. Yingxi Lin; Addgene plasmid # 63931; http://n2t.net/addgene:63931; RRID: Addgene_63931). Overexpression of Dnmt3a2 or a Control was achieved by using published and validated constructs^34^. For the expression of the catalytic-mutant form of Dnmt3a2, the previously validated inactive catalytic domain was kindly provided by Prof. Dr. Albert Jeltsch (University of Stuttgart, Germany) ^19^ and exchanged using Gibson Cloning with the wild-type one. For the chemogenetic inhibition of cortical engram neurons, the Cre-recombinase (CreER^T2^) under the cFos promoter was kindly provided and validated by Dr. Michel van den Oever (Vrije Universiteit Amsterdam)^4^. The mCherry control construct (pAAV-hSyn-DIO-mCherry) and the inhibitory DREADD construct (h4MDi: pAAV-hSyn-DIO-hM4D(Gi)-mCherry) was a gift from Bryan Roth (Addgene plasmid # 50459; http://n2t.net/addgene:50459; RRID: Addgene_50459): Addgene plasmid # 44362; http://n2t.net/addgene:44362; RRID: Addgene_44362). The TRAP-based viral vectors were used as previously described^4^.

### Stereotaxic delivery

rAAVs were injected at specific coordinates relative to Bregma: dHPC: AP: −2 mm; ML: ±1.4 mm; DV: −1.3 (650 nl) and DV: –2.1 (300nl); ACC: AP: 0.5 mm; ML: ±0.3 mm; DV: −1.5 (650nl). rAAVs were injected at a speed of 135 nl/min using a 33-G Nanofil needle (WPI, Sarasota, FL, USA). Before and after injections at each individual spot, the needle was left in place for 5 minutes. All viruses (except TRAP system) were used at a final working concentration of 10^12^ viral particles/mL. For the viral TRAP system, a virus mixture of rAAV-Fos-CreERT2 (titre: 1.4×10^12^ viral particles/ml) and the Cre-dependent rAAV (titre: 5.0-6.0 ×10^12^) were injected in a ratio of 1:9.

### Contextual Fear Conditioning

Before behavioural training, mice were habituated to the experimenter (by gentle handling for 2 minutes) and testing environment for 3 days. Different cohorts of mice were used to assess recent (24 hours) or remote (2 weeks) memory. In experiments involving Dnmt3a2 overexpression or engram dynamics of short-lasting fear memory, mice were allowed to explore a conditioning chamber (23 × 23 × 35 cm, TSE, Bad Homburg, Germany) for 148 seconds before receiving a 0.2mA foot shock for 2 seconds, followed by a 30-second retention period before returning to their home cage. For studying engram dynamics of long-lasting fear memory, mice underwent a training session consisting of 148 seconds of exploration followed by three 0.7mA foot shocks for 2 seconds each, spaced by 148-second intervals, and a 60-second retention period. In loss-of-function studies (Dnmt3a2 knockdown), to prevent possible ceiling effects, mice were allowed to explore the conditioning chamber for 148 seconds before receiving a 0.7mA foot shock for 2 seconds, followed by a 30-second retention period before returning to their home cage. For chemogenetic inhibition of cortical engram neurons (viral TRAP system), mice received intraperitoneally 4-Hydroxytamoxifen (4-OHT) 2 hours after contextual fear conditioning and Clozapine N-oxide (CNO) 30 minutes before remote fear testing. Testing sessions involved exposing the animals to the conditioning chamber for 5 minutes. In the altered context experiment, the testing chamber differed significantly from the original chamber in terms of location, scent (lemon detergent instead of 70% ethanol), floor material (white plastic instead of a metal grid), shape (triangle instead of square), and light intensity. Freezing behaviour was manually scored by an experimenter blind to the virus identity.

### qRT-PCR primer design

The quantitative reverse-transcription PCR (qRT-PCR) primers were designed with Primer3 (https://primer3.ut.ee/) using the RefSeq curated annotation along with the GRCm38/mm10 mouse genome assembly. The specificity and amplicon product size of the primers were verified by BLAST search and *in silico* PCR (UCSC Genome Browser, mm10). Primer pair efficiencies and product melting curves were validated by qRT-PCR prior to use. The following primers were used: Actin: 5’Primer: TATCCTGACCCTGAAGTACC, 3’Primer: CTCGGTGAGCAGCACAGGG; Calb1: 5’Primer: GCGAGGAATTCATGAAGACTTGG, 3’Primer: TGTCAGTTCCAGCTTTCCGT; Fhad1: 5’Primer: AGACGAAAATGATCCTGACGG, 3’Primer: CTAGGCTCACGATGGTCTGC; Nfix: 5’Primer: TGTCCAGCCACATCACATTG, 3’Primer: TGGAAACTTAAGTGCCCGTTG; Nt3: 5’Primer: GCACAATACAGCTGGTCGTC, 3’Primer: TGGCAGTCACAAGCTCTG; Pcdh10: 5’Primer: TCGCGAGCAAATCTGTAAGC, 3’Primer: AGGAGGGAGGGTTGTCATTG; Plxcn1: 5’Primer: GCTGGGAAGGAGGTGAGAAG, 3’Primer: GTGCACCTTTGTAACGGGAG

### qRT-PCR

dHPC of mice was quickly dissected and RNA was extracted using the RNeasy Plus Mini Kit (Qiagen, Hilden, Germany) with extra DNase I digestion on the column, following the manufacturer’s instructions. RNA was transcribed into cDNA using the High-Capacity cDNA reverse-transcription kit (Applied Biosystems, Foster City, CA, USA). qRT-PCR was performed on Step One Plus Real-Time PCR System (Applied Biosystems, Foster City, CA, USA) using the Power SYBR Green PCR Master Mix (Applied Biosystems) assays as technical triplicates using 0.5 μM of each primer with 2 μL of diluted cDNA (about 1.25 ng) in each reaction. Thermal cycling was done with the following settings: a 10-minute incubation at 95 °C, 40 cycles of 10 seconds each at 95 °C, 60 °C, and 72 °C, followed by a 15-second incubation at 95 °C. Melt curves were generated by heating from 60 °C to 90 °C at a ramp rate of 0.6 °C/min. Relative expression levels of each target transcript were determined by the ΔΔCt method using beta-Actin mRNA levels as a reference^35^.

### Bulk RNA Sequencing

Total RNA from mouse hippocampal cultures was isolated as above described and 500 ng of total RNA was used for bulk RNA-sequencing. Data processing was performed with R (version 3.6.3) and bioconductor (version 3.9) in Rstudio (version 1.1.463). Quality control of clean sequencing reads was performed using FastQC (Babraham Bioinformatics). Low-quality reads were removed using trim_galore (version 0.6.4). The resulting reads were aligned to the mouse genome version GRCm38.p6 and counted using kallisto version 0.46.1^36^. The count data were transformed to log2-counts per million (logCPM) using the voom-function from the limma package^37^. Differential expression analysis was performed using the limma package in R. A false positive rate of α= 0.05 with FDR correction was taken as the level of significance. Volcano plots were created using ggplot2 package (version 2.2.1)^38^. For enrichment analysis, ShinyGO online platform was used^39^. Pathway size of 20 was used and GO terms were selected by FDR.

### Fluorescent staining

Mice were anesthetised with Narcoren (2 μl/g body weight; Merial GmbH) and perfused intracardially with ice-cold PBS followed by ice-cold 4% paraformaldehyde (PFA) (Sigma-Aldrich, Munich, Germany). Brains were collected and post-fixed in 4 % PFA over night at 4 °C. To achieve cryoprotection, brains were placed in a 30 % sucrose solution in PBS with 0.01 % Thiomersal (Carl Roth) and cut at 30 μm thickness using a Leica CM1950 Cryostat. For fluorescent staining, slices were washed in PBS and blocked in 8 % normal goat serum with 0.3 % Triton X-100 in PBS for 60 min at room temperature and washed once with PBS. Primary antibody (mouse-anti-HA, Covance (1:1000); rabbit-anti-c-fos, Cell Signaling (1:2000)) were diluted in 2 % normal goat serum with 0.3 % Triton X-100 in PBS and incubated with slices over night at 4 °C mildly shaking. After three washing steps with PBS for 5 min, slices were incubated with secondary antibody (diluted 1:500) in 2 % normal goat serum with 0.3 % Triton X-100 in PBS for 90 min in the dark at room temperature and washed three times for 5 min with PBS. Finally, slices were incubated in Hoechst 33258 (2μg/ml, Serva, Heidelberg, Germany) for 5 min and mounted on glass slides.

### Image acquisition and analysis

Image acquisition was done using a 20x oil objective on a TCS SP8 confocal microscope (Leica Microsystems, Oberkochen, Germany) and the Leica LAS X LS software. 10% overlap ratio was set to stitch individual tiles of a bigger image. Image analysis was done using Fiji. Background fluorescence was subtracted. For overlap analysis, a threshold was defined, and particles analysed. Only particles larger than 30-pixel units and above the threshold were included in the analysis. The same threshold of intensity was applied to all images to identify cells positive for GFP or cFos. Cell-counter plugin of Fiji was used to count the identified particles in each channel independently. A particle was designated as “overlapping” when it was positively identified in both channels. The total number of cells (Hoechst^+^ cells) was calculated using a formula based on the total area of the DG, CA1 or ACC which was developed and generated using cell count and area from 15-20 mice (DG: R2=0.91; CA1 R2=0.96, ACC: R2=0.97). Throughout the entire study, the image analysis was performed on at least 3 animals per condition and at least 3 brain slices per animal. The final value for each animal represents the average of its brain slices. The reactivation rate ((GFP^+^cFos^+^)/(GFP^+^) x100) was calculated as previously described^15,40^

### Statistical analysis

Each data set was subjected to a normality test prior to further comparisons (Shapiro-Wilk normality test; alpha = 0.05). For normally distributed data sets, a two-tailed unpaired Student’s t-test was performed to compare two groups. If more than two groups were analysed simultaneously, a one-way ANOVA was used followed by appropriate multiple comparison *post hoc* tests to control for multiple comparisons as specified. The sample size was determined based on similar experiments carried out in the past and the literature. All plotted data represent mean ± SEM. Statistical analysis was performed using GraphPad Prism 7 (GraphPad Software).

## Acknowledgements

This work was supported by the Deutsche Forschungsgemeinschaft (DFG) [grant numbers OL 437/2, OL 437/3, OL 437/4 and OL 437/7 to A.M.M.O.], the Chica and Heinz Schaller Foundation [fellowship and research award to A.M.M.O.], and the Joachim Herz Stiftung [Add-on Fellowships for Interdisciplinary Life Science to J.K.].

## Author Contributions

JK, AMMO designed research; JK, SL & CPB performed experiments and analysed data; CS performed RNA Sequencing analysis; JK & AMMO interpreted data; JK & AMMO wrote the paper. All authors edited the manuscript.

---

**Extended Data Fig 1.**
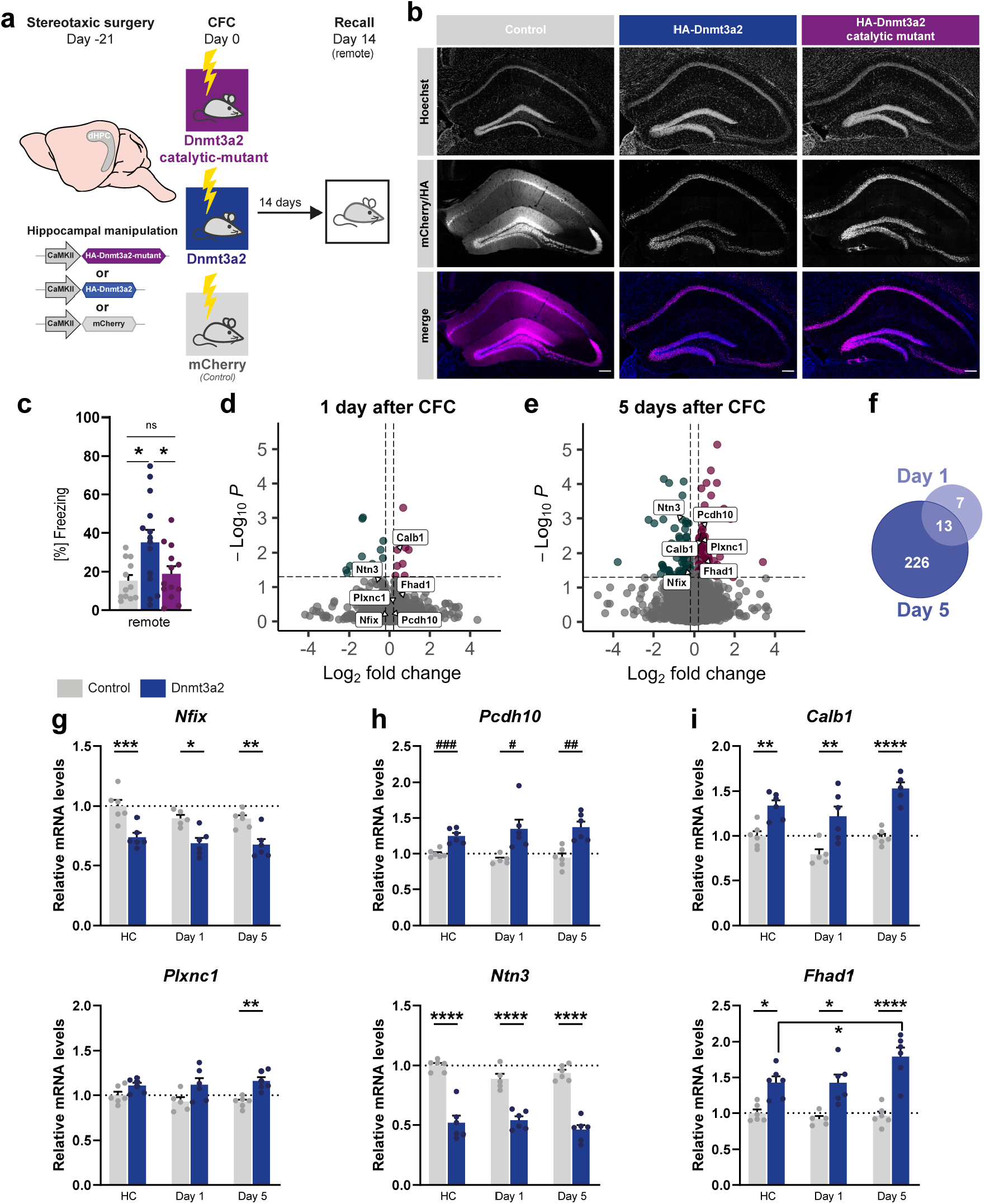
Dnmt3a2 regulates transcription of genes involved in synaptic transmission. (a) Schematic representation of experimental design. Stereotaxic injection of rAAV to overexpress a control, Dnmt3a2 or a catalytic-inactive Dnmt3a2 in dHPC. Three weeks after surgery, mice were trained in CFC (1x 0.2 mA shock) their memory was tested in a recall session 14 days after. (b) Representative images of dHPC of mice injected with Control (mCherry), Dnmt3a2 overexpression construct or catalytic-inactive variant. Scale bar: 100μm. (c) Remote contextual fear memory of mice injected with rAAVs overexpressing mCherry (n=12), Dnmt3a2 (n=14) or catalytic-inactive form of Dnmt3a2 (n=13). *p<0.05, ns: not significant by one-way ANOVA test followed by Sidak’s multiple comparisons test. (d, e) Volcano plot of DEGs (d) 1 day or (e) 5 days after CFC training. (f) Venn-Diagram of DEGs 1 day and 5 days after CFC training. (g, h, i) qRT-PCR analysis of genes in dHPC of mice injected with rAAVs to erexpress mCherry (n=5-6) or overexpress Dnmt3a2 (n=6) that either stayed in their home cage (HC) or underwent CFC and were sacrificed 1 day or 5 days after. Expression levels were normalised to the mCherry HC group (dashed line). *p<0.05, **p≤0.01, ***p≤0.001, ****p≤0.0001 by one-way ANOVA test followed by Sidak’s multiple comparisons test; ^#^p<0.05, ^##^p≤0.01, ^###^p≤0.001 by two-tailed, unpaired t-test. Error bars represents S.E.M.

**Extended Data Fig 2.**
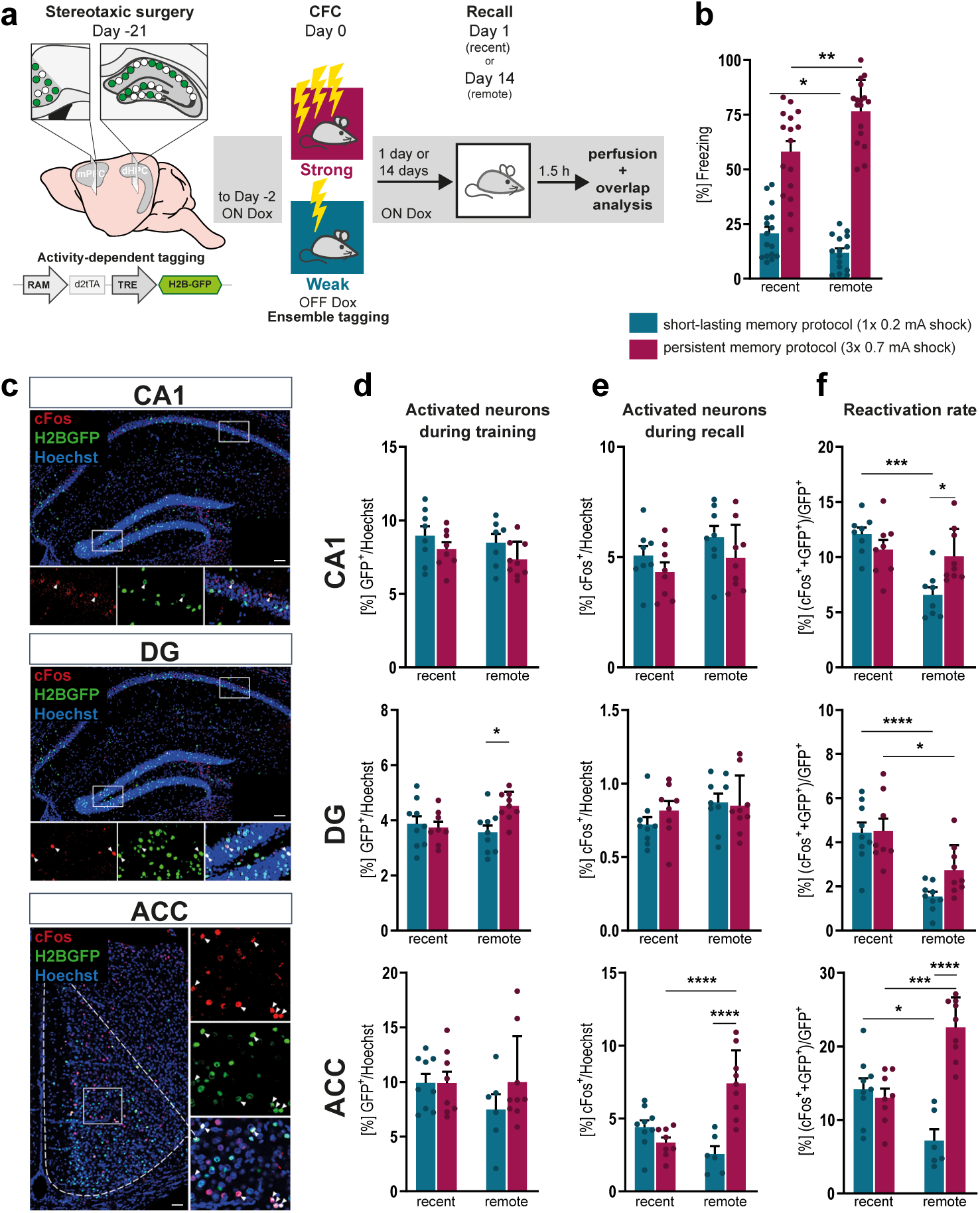
Engram reactivation dynamics in different fear memory paradigms. (a) Schematic representation of experimental design. Stereotaxic injection of rAAV to tag CA1, DG and ACC engram cells. Three weeks after surgery, mice were trained in CFC (1x 0.2 mA shock or 3x 0.7 mA shock) and their memory was tested in a recall session 1 day or 14 days after. 1.5h post recall, mice were sacrificed for overlap analysis. (b) Recent or remote contextual fear memory of mice (1x 0.2 mA shock: n= 16(recent); n=15(remote) or 3x 0.7 mA shock: n=16(recent); n=16(remote)). *p<0.05, **p≤0.01 by two-tailed, unpaired t-test. (c) Representative images of overlap analysis of CA1, DG and ACC. H2BGFP (green) indicates the neuronal population activated by training; cFos (red) represents the recall-activated neurons. (d, e, f) Quantitative image analysis in CA1, DG and ACC region of (d) activated neurons during training assessed by GFP signal, (e) activated neurons during recall identified by endogenous cFos labelling, (f) reactivation rate (GFP^+^+cFos^+^ neurons in GFP^+^ population (n=6-9). *p<0.05, **p≤0.01, ***p≤0.001, ****p≤0.0001 by one-way ANOVA test followed by Sidak’s multiple comparisons test

